# MTB-LysB1:A Novel Endolysin Against Multidrug-Resistant *Mycobacterium tuberculosis*

**DOI:** 10.64898/2026.07.13.738107

**Authors:** Ritu Arora, Eniyan Kandasamy, Jyoti Rani, Amit Kumar Singh, Urmi Bajpai

## Abstract

The phenotypic plasticity, slow replication, and complex, hydrophobic cell envelope of *Mycobacterium tuberculosis* contribute to its successful survival as a pathogen and its drug tolerance. Consequently, the global threat of multidrug-resistant Tuberculosis (MDR-TB), coupled with lengthy and highly toxic treatment regimens, necessitates the development of innovative treatment solutions. Mycobacteriophages are natural viruses of mycobacteria that typically encode two endolysins, which cooperatively facilitate host cell lysis at the end of the lytic life cycle: LysA, a peptidoglycan hydrolase, and LysB, a lipolytic enzyme, targeting the mycolylarabinogalactan-peptidoglycan complex. Their precise and efficient lytic activity, along with their low propensity to induce resistance, make them, particularly LysBs, promising candidates for new treatment solutions.

In this study, we report MTB-LysB1, a novel LysB enzyme from an F1 sub-cluster mycobacteriophage isolated from our laboratory collection. While studying its structural features by comparing the modelled structure with representative mycobacteriophage LysB homologues, we found that the α/β-hydrolase fold and key motifs are conserved. Also, we identified putative membrane-interaction motifs that may play a role in LysB1’s cell permeation. Significantly, we found MTB-LysB1 to be active against both drug-susceptible and multidrug-resistant (MDR) *M. tuberculosis* strains at nanomolar concentrations, comparable to the well-characterised D29 LysB reference enzyme. Beyond its standalone activity, MTB-LysB1 exhibits an additive effect when combined with the TB drugs rifampicin and moxifloxacin, and co-administration reduces the drugs’ minimum inhibitory concentrations (MICs), which holds clinical significance. By structurally damaging the mycobacterial cell wall, the enzyme appears to act as a permeability enhancer for the chemotherapeutic drugs, thereby improving antibiotic efficacy. Collectively, our findings position the enzyme not only as a novel antimycobacterial agent but also provide a structural framework for its rational engineering as a promising next-generation adjunct to TB drug regimens.

**Highlights:**

1. A novel F1 sub-cluster phage-derived LysB is discovered and characterised using integrated computational, biochemical and microbiological methods.
2. AlphaFold2 modelling, molecular dynamics simulations and comparative structural analyses revealed an α/β-hydrolase fold with conserved catalytic and membrane-interaction features.
3. The enzyme exhibited high esterase activity, thermal stability and potent lytic activity against *Mycobacterium tuberculosis*.
4. An additive effect with TB drugs rifampicin and moxifloxacin highlights MTB-LysB1’s potential as an adjunct therapeutic.

## Introduction

Endolysins are the bacteriophage-encoded lytic enzymes that assist in the release of progeny phages by breaching the host bacterium’s cell wall. These enzymes have evolved to lyse host cells from within. However, some of the endolysins also exhibit properties that cause cell lysis when applied externally. Because of their remarkable ability to selectively target and damage bacterial cell integrity, endolysins are being investigated asa new class of protein antibiotics in the wake of the antimicrobial resistancecrisis [1–7]. Endolysins offer several advantages over conventional antibiotics, most notably their high specificity, in contrast to the broad-spectrum activity of many antibiotics,andtheir markedly low tendency for target bacteria to develop resistance [1,8]. In addition, endolysins exhibit rapid bacteriolytic activity,synergistic/additive effects when combined with existing antibiotics, and efficacy against bacterial biofilms [9–12]. In several genera, heterologous expression of recombinant lysins has been achieved, and their antibacterial activity has been well documented. Examples include *S. aureus* [13,14], *S. pneumoniae* (including β-lactam-resistant strains) [15], and *C. difficile* [16]. Collectively, these properties make endolysins highly promising adjuncts to existing therapies for the treatment of drug-resistant bacterial infections [4,7,11,17,18,19].

Most mycobacteriophages encoded two distinct endolysins: LysA and LysB. The two-enzyme system is an adaptation to the complex mycobacterial cell envelope, which consists of a cytoplasmic membrane surrounded by a cell wall composed of a thin peptidoglycan layer covalently linked to the arabinogalactan (AG). The AG layer, in turn, is esterified to long-chain mycolic acids (MA), forming the mycolyl-arabinogalactan peptidoglycan complex (mAGP). To exit the host, LysA acts upon the peptidoglycan layer, and LysB, acting as a mycolyl-arabinogalactan esterase, cleaves the ester linkage between the mycolic acids and the arabinogalactan, effectively disrupting the outer mycolic acid barrier and releasing free mycolic acids [20]. Figure 1 describes both endolysins’ site of action. LysB offers greater therapeutic potential and specificity than LysA because it directly targets the mycomembrane, the outermost defence characteristic of *Mycobacterium*. By compromising the structural integrity of the hydrophobic envelope, LysB can significantly enhance the pathogen’s susceptibility to synergistic drug treatments.

**Figure 1.**
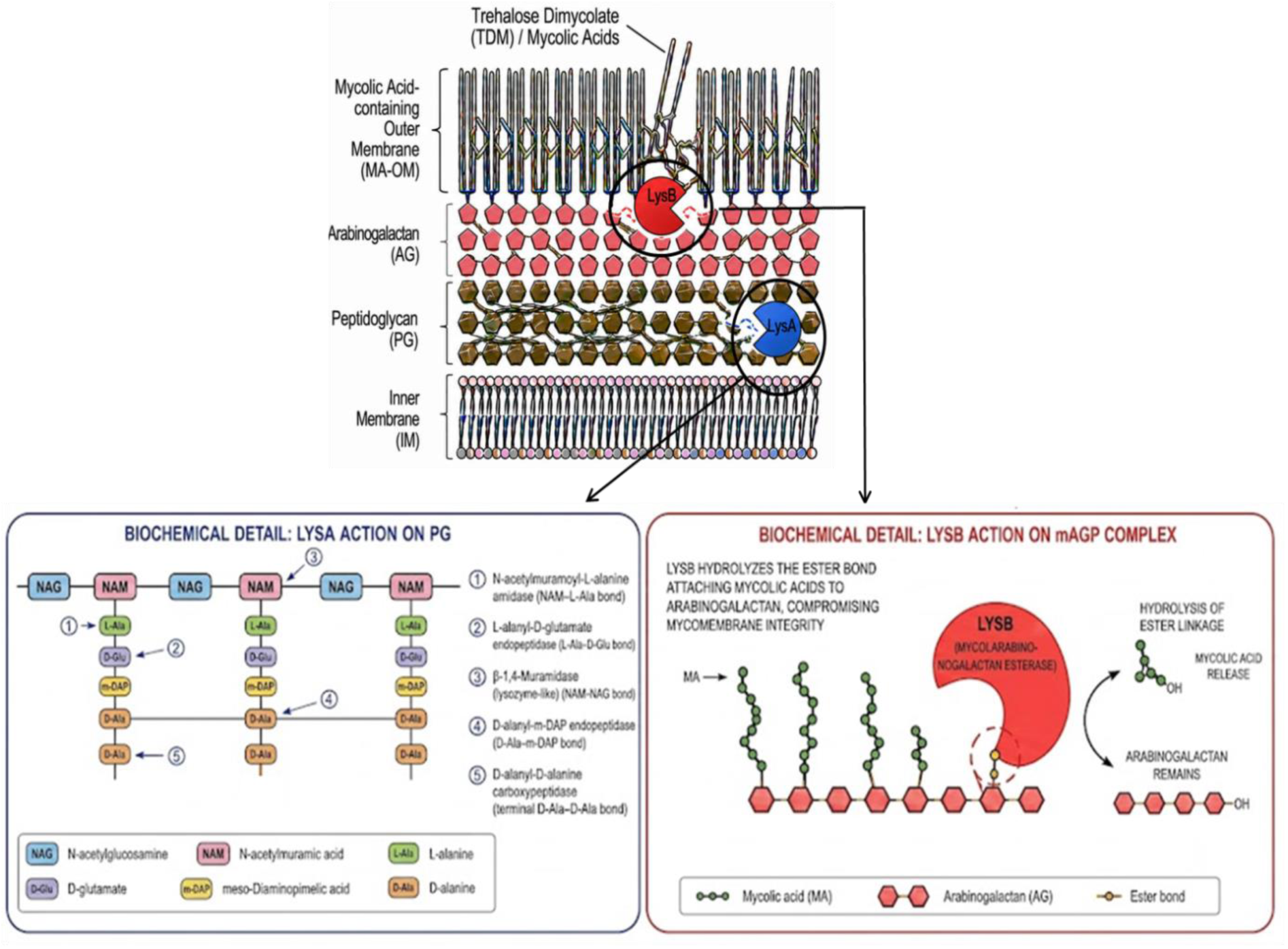
Schematic diagram showing the cleavage sites of endolysins on the mycobacterial cell envelope. LysA targets the peptidoglycan backbone, LysB targets the ester bond between MA and AG, weakening the outer membrane barrier. Adapted from Ceballos-Zúniga et al. [21]. The image was generated using Google Gemini.

In this study, we introduce a novel LysB ‘MTB-LysB1’ which we purified from an F1 sub-cluster mycobacteriophage from our laboratory phage collection. We have described its antimycobacterial activity and compared its structural features with those of two well-studied LysBs: D29 and Ms6. Based on the structural alignment with known LysB structures, we noted that MTB-LysB1 is similar to members of the α/β hydrolase family, which includes cutinases, esterases and lipases. Its catalytic triad is constituted of Ser169-Asp253-His318, where serine is also part of the [G-X-S-X-G] motif. While the positions of Ser and Asp residues in the catalytic triads are reported to be highly conserved, the position of the His residue shows a slight shift [22]. Further, to characterise the domain structureand compare it with D29 and Ms6 LysBs, we modelled the MTB-LysB1 structure using AlphaFold2, followed by MD simulation to analyse its stability and binding efficiency. Since the structure of Ms6 LysB (also derived from an F1 sub-cluster mycobacteriophage) isn’t available, its structure was also modelled. To estimate MTB-LysB1’s enzymatic (esterase) activity and stability, the pNP-release assay was performed, and to determine its anti-bacterial activity, three methods were used using *M. smegmatis* Mc^2^ 155 as the target: plate lysis, turbidity reduction, and log kill assays. Finally, anti-*M. tuberculosis* activity (against H37Rv and an MDR isolate) of MTB-LysB1 was assessed by plate lysis followed by a REMA assay, alone and in combination with TB drugs.

## Materials and Methods

### Computational Analysis

To assess sequence homology and domain architecture, the MTB-LysB1 protein sequence was analysed using BLASTp [23], InterProScan [24] and HMMER [25]. ClustalOmega (1.2.4) [26] was used for multiple sequence alignment.

#### Structural Modelling and Comparative Analysis

The three-dimensional structure of the uncharacterised MTB-LysB1 protein was predicted *de novo* using AlphaFold2 [27]. For comparative purposes, the structures of RitSun [12], Ms6 [28], and TM4 [29] LysBs were also modelled using the same protocol to maintain methodological consistency. Model quality was evaluated based on per-residue confidence scores (pLDDT) and overall Template Modelling (TM) scores, which reflect local reliability and global fold accuracy, respectively.

#### Molecular Dynamics (MD) Simulation

To assess the structural stability of the predicted MTB-LysB1 structure under physiological conditions, all-atom MD simulations were performed using Desmond (Schrödinger LLCat ICGEB, New Delhi). The protein was placed in an orthorhombic water box using the TIP3P solvent model and neutralised with appropriate counter ions. System energy minimisation was followed by equilibration under NVT and NPT ensembles at 300 K and 1 atm pressure. Production runs were carried out for 100 ns, with trajectory analyses conducted to calculate Root-Mean-Square Deviation (RMSD), Root-Mean-Square Fluctuation (RMSF), radius of gyration (Rg), and secondary structure stability. These analyses provided insights into the conformational flexibility, domain stability, and overall resilience of MTB-LysB1 under dynamic conditions.

#### Structural Validation and Alignment

To evaluate structural conservation and similarity, the predicted MTB-LysB1 model was pairwise aligned with the crystal structure of D29 LysB (PDB ID: 3HC7) and with the predicted structures of Ms6, TM4, and RitSun LysBs from this study, using the TM-align algorithm. The resulting TM-scores and Root Mean Square Deviations (RMSD) were calculated to quantify the extent of structural similarity. Superimposed structures were visualised in PyMOL, enabling detailed inspection of conserved folds and domain orientations across LysB homologues.

#### Binding Cavity and Functional Site Analysis

The potential ligand-binding and catalytic cavities of MTB-LysB1, D29, Ms6, TM4, and RitSun LysBs were predicted using the CavityPlus server. The top-ranked cavities were analysed based on their geometry, size, and residue composition. Spatial overlap of the identified pockets was examined in PyMOL to assess conservation of the putative active site. Key residues forming the catalytic triad were compared across homologues to identify conserved functional motifs relevant to enzymatic activity.

#### Analysis of Membrane Interaction Motifs

Putative hydrophobic and amphipathic helices potentially involved in membrane association were predicted from the MTB-LysB1 primary sequence using the HeliQuest server. These motifs were mapped onto the modelled structure and analysed in relation to the identified binding cavity to infer the protein’s orientation relative to the membrane. Comparative visualisation with D29 and other homologues was used to determine whether the membrane-interacting regions are structurally conserved.

### Bacterial Strains, Plasmids and Culture Conditions

*Escherichia coli* (*E. coli*) strain DH5α was used for cloning and plasmid preparation, and *E. coli* BL21 (DE3) was used for protein overexpression. pET28a vector was used for cloning and expression of the MTB-LysB1 gene. Both *E. coli* strains were cultured in Luria-Bertani (LB) broth with shaking at 200 rpm or on LB agar plates. Kanamycin (50 µg/mL) was used as a selection marker.

### Reagents and Enzymes

Phusion High-Fidelity DNA Polymerase,T4 DNA ligase and Restriction enzymes were purchased from New England Biolabs (USA). RNase A was sourced from Bangalore Genei (India), and analytical-grade chemicals were purchased from Sigma-Aldrich. DNA and protein molecular weight markers were obtained from Thermo Fischer Scientific (USA). Ni-NTA agarose resin and nucleic acid purification kits were obtained from Qiagen (Germany). Phenylmethylsulfonylfluoride (PMSF), antibiotics and additional reagents were supplied by HiMedia Laboratories (India). Primary and secondary antibodies were purchased from Santa Cruz Biotechnology (USA).

### Expression of Recombinant MTB-LysB1

The MTB-LysB1 gene induction conditions included0.1 mM IPTG and incubation at 16°C for 16 h under shaking conditions (200 rpm). Following induction, cells were lysed by sonication at 90% amplitude with 30-sec on/off pulses for 10cycles. The lysate was clarified by centrifugation at 5000 × g for 45 min at 4°C, and both soluble and insoluble fractions were analysed to evaluate recombinant protein expression.

### Purification and Western Blot Analysis

Soluble recombinant MTB-LysB1 was purified by Ni-NTA affinity chromatography as described previously [12,30]. The protein was allowed to bind to Ni-NTA agarose for 2 h at 4°C. The column was sequentially washed with increasing concentrations of imidazole, and the protein was eluted at 250 mM. Eluted fractions were analysed by 12% SDS-PAGE and dialysed against a buffer (25 mM Tris, 150 mM NaCl), adjusted to pH 7.9. For Western blotting, the protein was transferred to a PVDF membrane at 90 V for 2 h. Subsequently, the membrane was blocked overnight at 4°C with 3% BSA. The His-tagged MTB-LysB1 was detected by sequential incubation with a 1:2000 dilution of mouse anti-His antiserum (Santacruz Biotech, USA) for 4 h, followed by a 1:10,000 dilution of anti-mouse IgG HRP-conjugated secondary antibody (45 min). Washing steps (PBS/PBS-T) were performed between antibody incubations. The blot was developed using 3,3-diaminobenzidine (DAB) and H_2_O_2_ as the substrate [12,30,31].

### Biochemical Assay

The esterase activity of MTB-LysB1 was quantified using the pNP-release assay [17]. p-Nitrophenol butyrate (pNPB) was used as the substrate, and the release of p-Nitrophenol (pNP) was measured by incubating varying concentrations of MTB-LysB1 (0.15–3μM) with 10 mM pNPB in 25 mM Tris buffer (pH 7.2) at 37°C for 15 min. The released pNP was quantified by measuring absorbance at 410 nm. The reaction without the enzyme served as the control. The experiment was performed three times in triplicate. The A_410_ values are reported as mean ± SD, and the specific activity was calculated using the PNP standard curve. To evaluate thermal stability, the pNP-release assay was performed at 45°C, 55°C, and 65°C. The release of pNP was measured after 15 min of incubation at each temperature. The absorbance values were calculated as mean±SD.

### Activity of MTB-LysB1 against *M. smegmatis* Mc^2^ 155

#### Qualitative Assays

All growth studies used Middlebrook 7H9 or 7H10 media containing glycerol (0.4%), carbenicillin (50 μg/ml), and Tween-80 (0.05%).

##### Lytic Effect of MTB-LysB1

To determine MTB-LysB1’s lytic effect on external application, the plate-lysis method was used. Briefly, MTB-LysB1 was spotted on an *M. smegmatis* Mc^2^ 155 lawn on 7H10 medium, and the plates were incubated at 37°C for approximately 48 h to observe a zone of clearance. To estimate the growth-inhibitory effect, *M. smegmatis* Mc^2^ 155 cells were treated with MTB-LysB1 at different concentrations for a fixed 4 h. The untreated and treated cells at 0 h and 4 h were serially diluted and spotted on 7H10 plates containing 0.05% Tween-80.

##### FESEM Analysis of LysB-treated M. smegmatis

Field Emission Scanning Electron Microscopy (FESEM) analysis of MTB-LysB1-treated *M. smegmatis* Mc^2^ 155 cells was performed as previously described [12]. Bacterial cells were treated with MTB-LysB1(2.5μM) for 4 h. The images were acquired at the Indian Institute of Technology, New Delhi, using a TESCAN instrument under 15,000 x magnification.

#### QuantitativeAssays

##### Turbidity Reduction Method (TRM)

TRM studies were conducted as previously described by Arora et al. [12]. Briefly, *M. smegmatis* Mc^2^ 155 cells, grown in 7H9 medium, were adjusted to an OD_600_ of 0.3 and treated with MTB-LysB1 (1 μM). After incubation at 37°C for 24 h at 200 rpm, OD_600_ was measured. *M. smegmatis* cells incubated without the enzyme were used as a control.

##### Log Kill Assay

To estimate log-killing, we followed the protocol developed by Grover et al. [32] with minor modifications. Briefly, *M. smegmatis* Mc^2^ 155 cells were diluted to an OD_600_ of 0.1 and treated with different concentrations of MTB-LysB1 (0.5 µM, 1 µM, 2.5 µM, and 5 µM) in a 96-well flat-bottomed plate and incubated at 37°C and 220 rpm for 24 h. The final volume per well was kept at 200 μl. Post-incubation, cells were serially diluted in PBS (pH 7.4) containing 0.05% Tween-80 and plated on 7H10 agar. The plates were incubated at 37°C until colonies appeared, and the CFU/ml was calculated. PBS-treated cells were used as a negative control.

#### Activity of MTB-LysB1against *M. tuberculosis*

Experiments involving *M. tuberculosis* H37Rv and the multidrug-resistant (MDR) strain were conducted at the Experimental Animal Facility of theNational JALMA Institute for Leprosy & Other Mycobacterial Diseases, Agra, India.The plate lysis method was used to study the effect of MTB-LysB1 on *M. tuberculosis* H37Rv and MDR *M. tuberculosis*, as previously described for *M. smegmatis*.

##### Checkerboard Assay with TB drugs

To quantitatively assess the activity of MTB-LysB1 against both strains (H37Rv and MDR), the Resazurin Microtiter Assay (REMA) was performed with adaptations from Palomino et al. [33] and Singh et al. [11]. To test against the H37Rv strain, two-fold serial dilutions of the enzyme and the TB drug were prepared in Middlebrook 7H9 broth supplemented with 10% albumin–dextrose–catalase (ADC) and 0.05% Tween-80, ranging from 0.025 to 0.0002 µM for MTB-LysB1 and 0.30 to 0.02 µM for rifampicin. For the MDR isolate, two-fold serial dilutions were performed over the range of 0.16 to 0.002 µM for MTB-LysB1 and 2.5 to 0.15 µM for moxifloxacin.

Actively growing *M. tuberculosis* cultures in mid-log phase (OD_600_∼0.6–0.8) were harvested by centrifugation (2,500 × g, 10 min), washed once, and resuspended in fresh 7H9–ADC–Tween-80. The bacterial suspension was adjusted to a 1.0 McFarland standard and diluted to approximately 1×10^5 CFU/mL, consistent with standardised REMA conditions for drug susceptibility testing; the same inoculum was used for minimum inhibitory concentration (MIC) determination with the MDR isolate. In microplates, 100 µL of each MTB-LysB1 dilution was dispensed, followed by 100 µL of the standardised inoculum, yielding a final volume of 200 µL per well. Peripheral wells were filled with 200 µL sterile distilled water to minimise edge evaporation during prolonged incubation. Rifampicin was included as a positive drug control for drug-susceptible H37Rv, and moxifloxacin was used as a positive control for the MDR isolate in line with prior REMA evaluations of second-line drugs. Growth controls contained bacteria without LysB or drugs, and sterility controls contained medium only. Plates were sealed and incubated at 37°C for 6–7 days, consistent with established REMA protocols for *M. tuberculosis*.

After incubation, 30 µL of resazurin solution (0.02% w/v in sterile water) was added to each well, and the plates were incubated for an additional 24 h at 37°C, in the dark. Wells retaining a blue/purple colour were interpreted as growth inhibition, whereas a change to pink indicated active bacterial metabolism. The MIC was defined as the lowest concentration of MTB-LysB1 that prevented a colour change from blue/purple to pink, following accepted REMA definitions and prior MIC determinations for D29 LysB [11]. All assays were performed in at least duplicate.

## Results and Discussion

### Structural Analysis of MTB-LysB1

MTB-LysB1 (333 aa) protein sequence analysis using InterProScan showed a N-terminal region (1-93 aa), the presence of an alpha/beta hydrolase fold (94-331 aa) and a C-terminal linker domain (260-324 aa) (Supplementary Figure S1). When aligned with the selected LysB homologues D29, Ms6, RitSun and TM4 LysB, the G-X-S-X-G pentapeptide and GXP motifs, which are essential for esterase activity [30], were found to be conserved in MTB-LysB1; G-Y-S-Q-G and GNP, respectively. The catalytic triad Ser-Asp-His, where Ser is part of the pentapeptide motif, is essential for the catalytic activity of the LysB enzymes. In MTB-LysB1, Ser (169) and Asp (253) were observed to be conserved, whereas His (318) was noted to shift by one base [22].

#### LysB Proteins Structure Prediction

We predicted the structures of MTB-LysB1, Ms6, TM4 and RitSun LysB using AlphaFold2. The modelled structures exhibited high confidence, with Template Modelling (TM) scores of 0.80, 0.81and 0.81, respectively, indicating reliable folding predictions and well-defined tertiary structures. However, the TM score of TM4 LysB was 0.67, indicating low structural confidence and distinct structural conservation (Figure 2 and Supplementary Figure S2).

**Figure 2.**
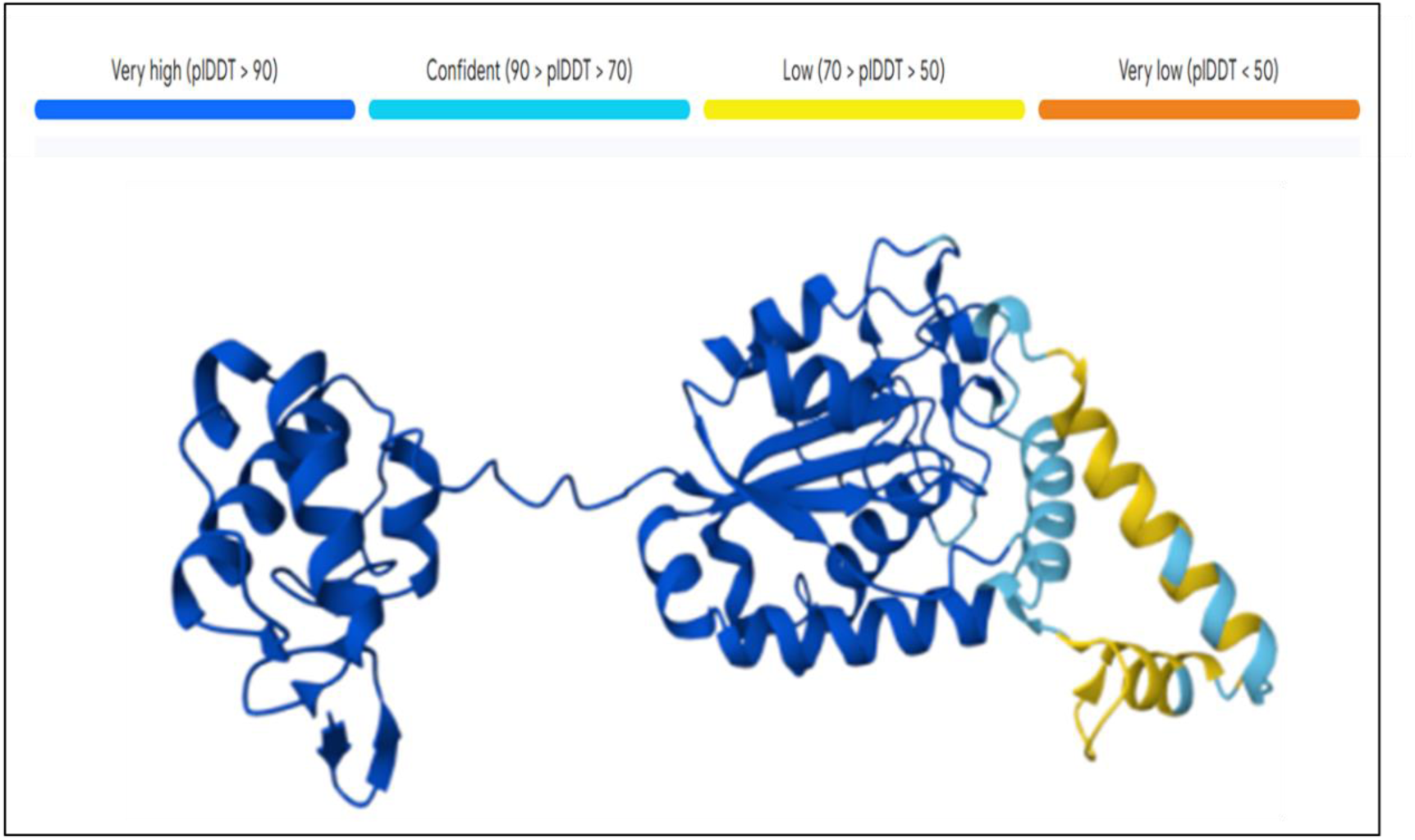
AlphaFold2 Modelled structures of MTB-LysB represented in cartoon format and coloured according to the confidence score. The protein exhibits a conserved central catalytic domain, predominantly composed of α-helices and β-sheets, connected to the terminal domain via a flexible linker region.

#### Molecular Dynamics (MD) Simulation

The dynamic behaviour and stability of MTB-LysB1 were evaluated through a 100 ns Desmond MD simulation under NPT conditions at 300K. RMSD analysis revealed that the protein backbone stabilised early in the simulation, fluctuating within 1-3 Å, which is indicative of equilibrium and the absence of major conformational changes (Figure 3A), suggesting that the protein maintains a stable overall fold during the simulation. RMSF analysis showed that loop regions and the N- and C-terminal residues exhibited the highest flexibility, whereas α-helices and β-strands remained relatively rigid, consistent with typical globular protein behaviour (Figure 3B). Monitoring of secondary structure elements (SSE) confirmed that the α-helices and β-strands were maintained throughout the trajectory (Figure 3C). The overall secondary structure composition remained stable, supporting the structural integrity of the predicted model. Overall, the MD simulation validated MTB-LysB1 as a structurally stable enzyme, suitable for functional analyses.

**Figure 3.**
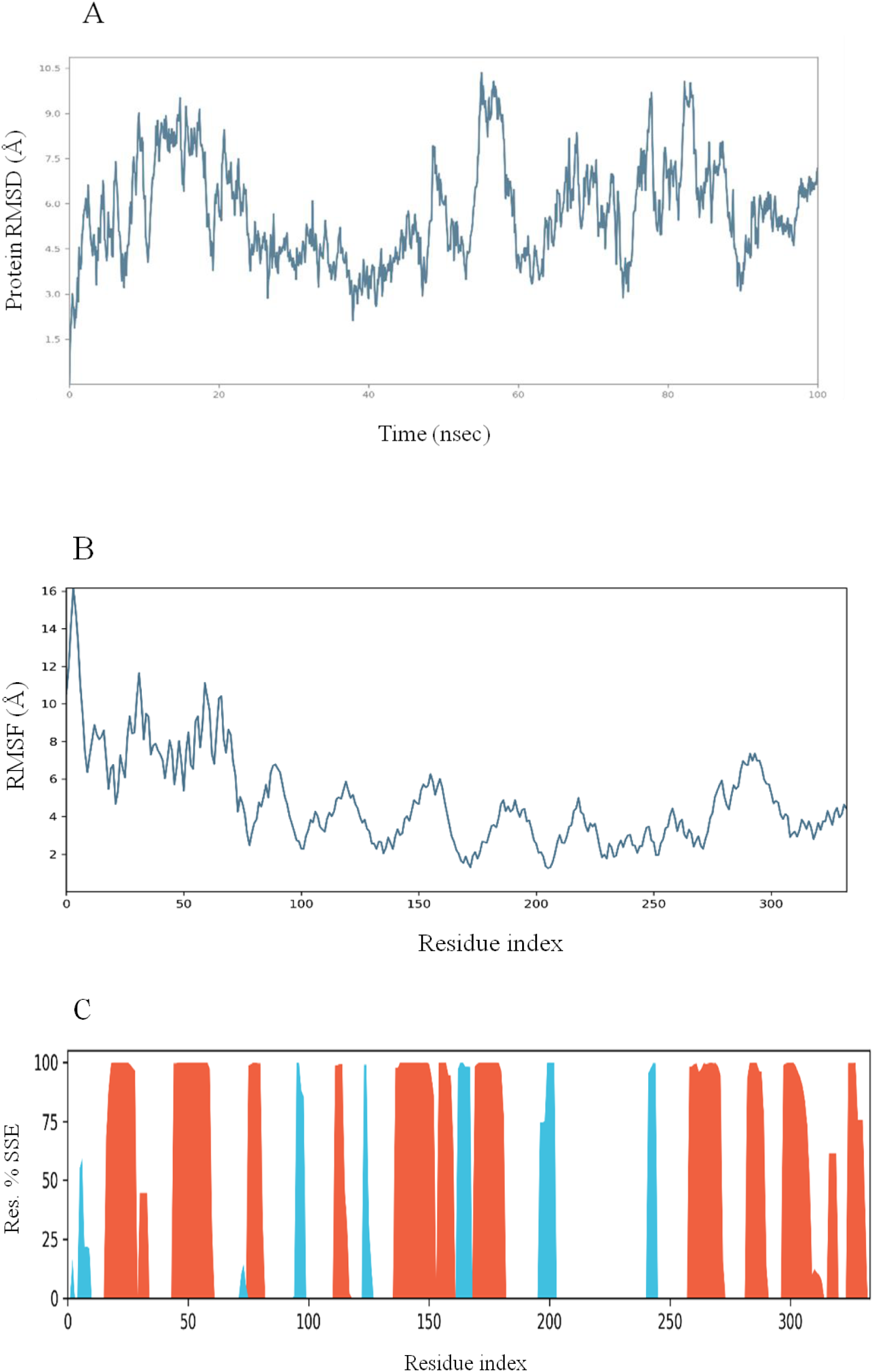
MD simulation: **(A)** Backbone RMSD shows structural stabilisation. **(B)** Residue-wise RMSF highlighting flexible loops and stable secondary structures. **(C)** SSE histogram demonstrating maintenance of α-helices and β-strands throughout the simulation.

#### Binding Cavity Characterisation of MTB-LysB1

To identify potential ligand-binding regions andassess the catalytic environment of MTB-LysB1, cavity analysis was performed using CavityPlus (Figure 4). Among the predicted pockets, Cavity 2, located within the protein’s central catalytic region, was found to be formed by residues previously identified as part of the conserved active-site pocket. Mapping of the cavity residues revealed a well-defined binding environment composed of hydrophobic, polar, and charged amino acids, supporting its potential role in substrate recognition and catalysis.

**Figure 4.**
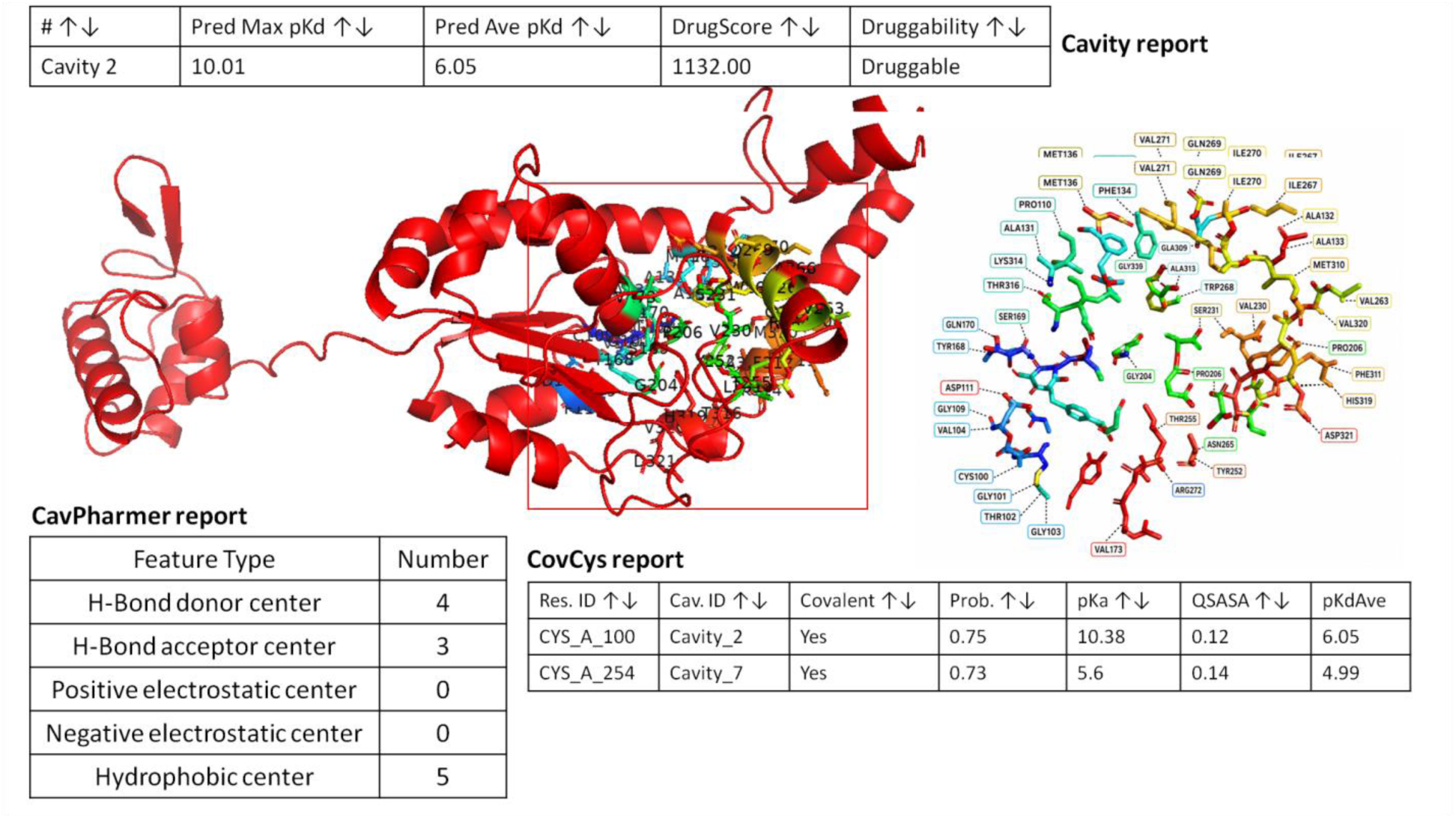
CavityPlus-based identification of the principal binding cavity (Cavity 2) in MTB-LysB1. The figure shows the cavity location, cavity-lining residues and CovCys-predicted reactive cysteine residues.

Pharmacophore analysis of the cavity identified four hydrogen-bond donor centres, three hydrogen-bond acceptors, and five hydrophobic centres, indicating multiple interaction hotspots that could facilitate substrate binding. Furthermore, CovCys analysis predicted Cys100 as a potential reactive cysteine residue within Cavity 2, with a probability score of 0.75 and a predicted pKa of 10.38. The location of this residue within the identified cavity suggests that it may contribute to ligand interactions. Together, these results indicate that Cavity 2 represents the primary functional pocket of MTB-LysB1 and provides a structural basis for its catalytic activity and ligand-binding properties.

#### Structural Alignment of Modelled LysB Proteins with the Reference Template (D29 LysB)

To evaluate the structural accuracy of the AlphaFold2-predicted models of MTB-LysB1 and three selected LysB homologues (Figure 2, Supplementary Figure S2), TM-align analysis was conducted using the crystal structure of D29 LysB (PDB ID: 3HC7) as a template (Figure 5). The analysis revealed that all four modelled proteins exhibited substantial structural similarity to the reference fold, despite moderate sequence divergence, supporting a common evolutionary origin and shared catalytic functions.

**Figure 5.**
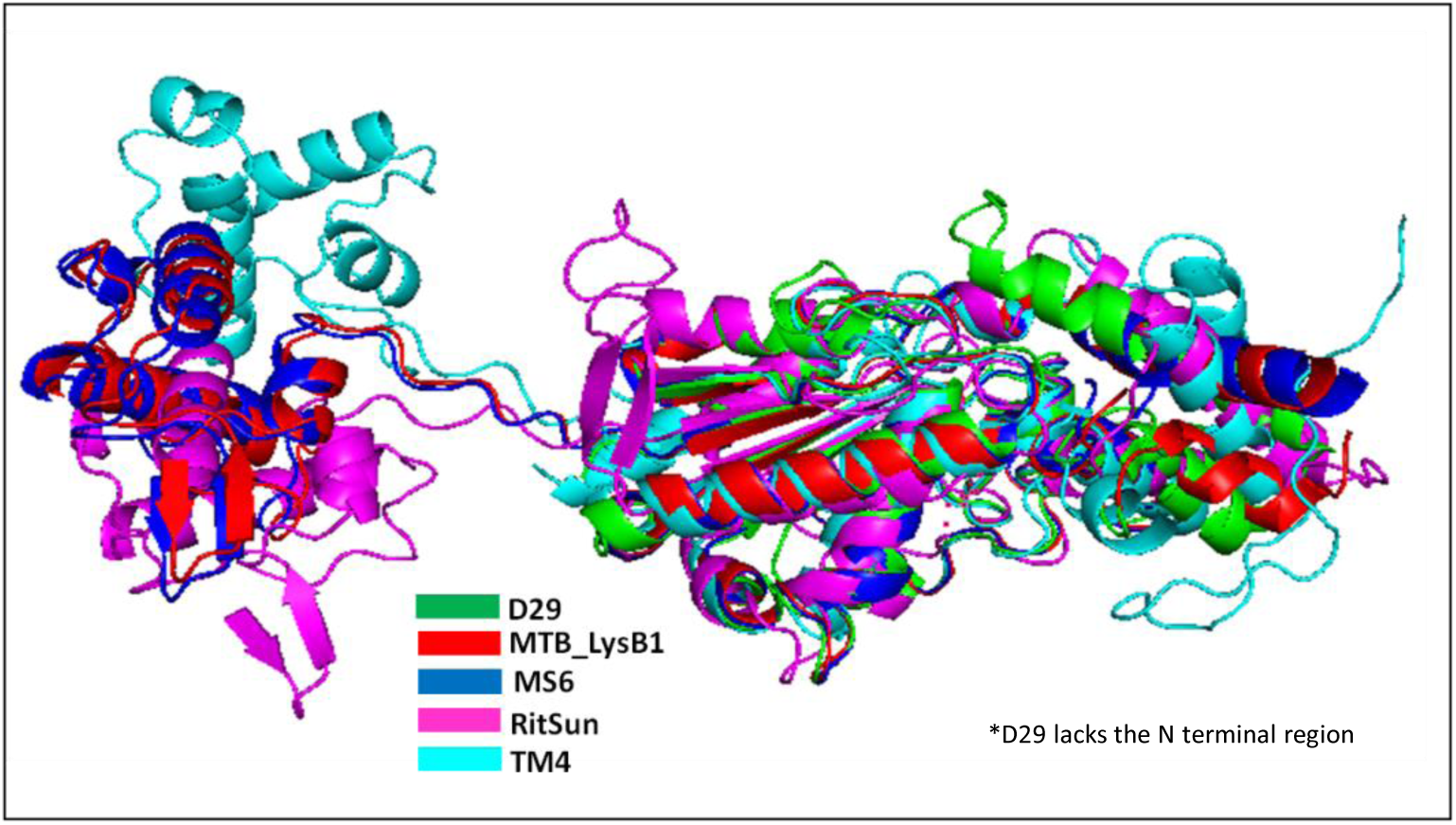
Superimposed protein structures of D29 (green), MTB-LysB1 (red), Ms6 (blue), RitSun (magenta), and TM4 (cyan) showing overall structural conservation with localised conformational differences, except for the lack of an N-terminal region in D29. The core α-helical and β-sheet regions remain largely conserved across the proteins, whereas variations are observed mainly in loop and terminal regions.

The MTB-LysB1 model aligned with 3HC7 over 214 residues, yielded a TM-score of 0.84 and an RMSD of 2.49Å, indicating strong structural conservation. Similarly, Ms6 LysB showed a TM-score of 0.82 (RMSD=2.52Å, 210 aligned residues), further confirming the reliability of its predicted tertiary structure. In comparison, RitSun LysB and TM4 LysB displayed somewhat lower TM-scores of 0.77 and 0.78, respectively, accompanied by higher RMSD values (3.43Å and 3.39Å) and fewer aligned residues (198 each) (Table 1). These differences suggest possible structural deviations arise from sequence divergence (22–27% identity) relative to 3HC7.

**Table 1.** Structural alignment of modelled LysBs with D29 LysB (PDB ID: 3HC7).

| Protein | Chain | RMSD | TM-score | Identity | Aligned Residues | Sequence Length | Modelled Residues |
| --- | --- | --- | --- | --- | --- | --- | --- |
| **3HC7(D29)** | A | - | - | - | - | 254 | 252 |
| MTB-LysB1.pdb | A | 2.49 | 0.84 | 38% | 214 | 333 | 333 |
| Ms6 LysB.pdb | A | 2.52 | 0.82 | 37% | 210 | 332 | 332 |
| RitSun LysB.pdb | A | 3.43 | 0.77 | 22% | 198 | 356 | 356 |
| TM4 LysB.pdb | A | 3.39 | 0.78 | 27% | 198 | 400 | 400 |

The structural superimposition (Figure 5) of five proteins further confirmed a highly conserved catalytic core composed of α-helices and β-sheets with structural variations primarily localised to the terminal and flexible loop regions. The prominent distinction is the absence of an N-terminal extension in D29 [34] that is consistently present alongside the catalytic esterase domain in the other homologues. Because the N-terminal region of Ms6 LysB is reported to bind to *M. smegmatis* and *M. tuberculosis* cells, suggesting a role in cell-envelope recognition and interaction [28,35], the conservation of this region in MTB-LysB1 and other proteins indicates equivalent mycobacterial cell-wall-binding capabilities.

Overall, the alignment results demonstrate that all modelled LysB proteins preserve the canonical α/β-hydrolase fold typical of the LysB family, and MTB-LysB1 combines the conserved catalytic architecture of LysB proteins with an additional N-terminal module. This strong structural conservation validates the AlphaFold2 predictions and supports the functional relevance of the modelled enzymes for subsequent comparative and dynamic analyses.

*Binding Cavity and Functional Site Analysis of* MTB-LysB1, Ms6 and D29 *LysB proteins*

Given the strong structural similarity ofMTB-LysB1, Ms6 and D29 among the five tested LysBs, further comparison in binding-cavity analysis was performed in these three LysB proteins to assess functional conservation. CavityPlus predictions revealed that MTB-LysB1, Ms6, and D29 share overlapping catalytic pockets with similar geometries and residue compositions. The conserved positioning of key catalytic triad residues across all homologues indicates shared enzymatic activity, substrate specificity, and membrane interaction potential, confirming functional equivalence within the LysB family (Figure 6).

**Figure 6.**
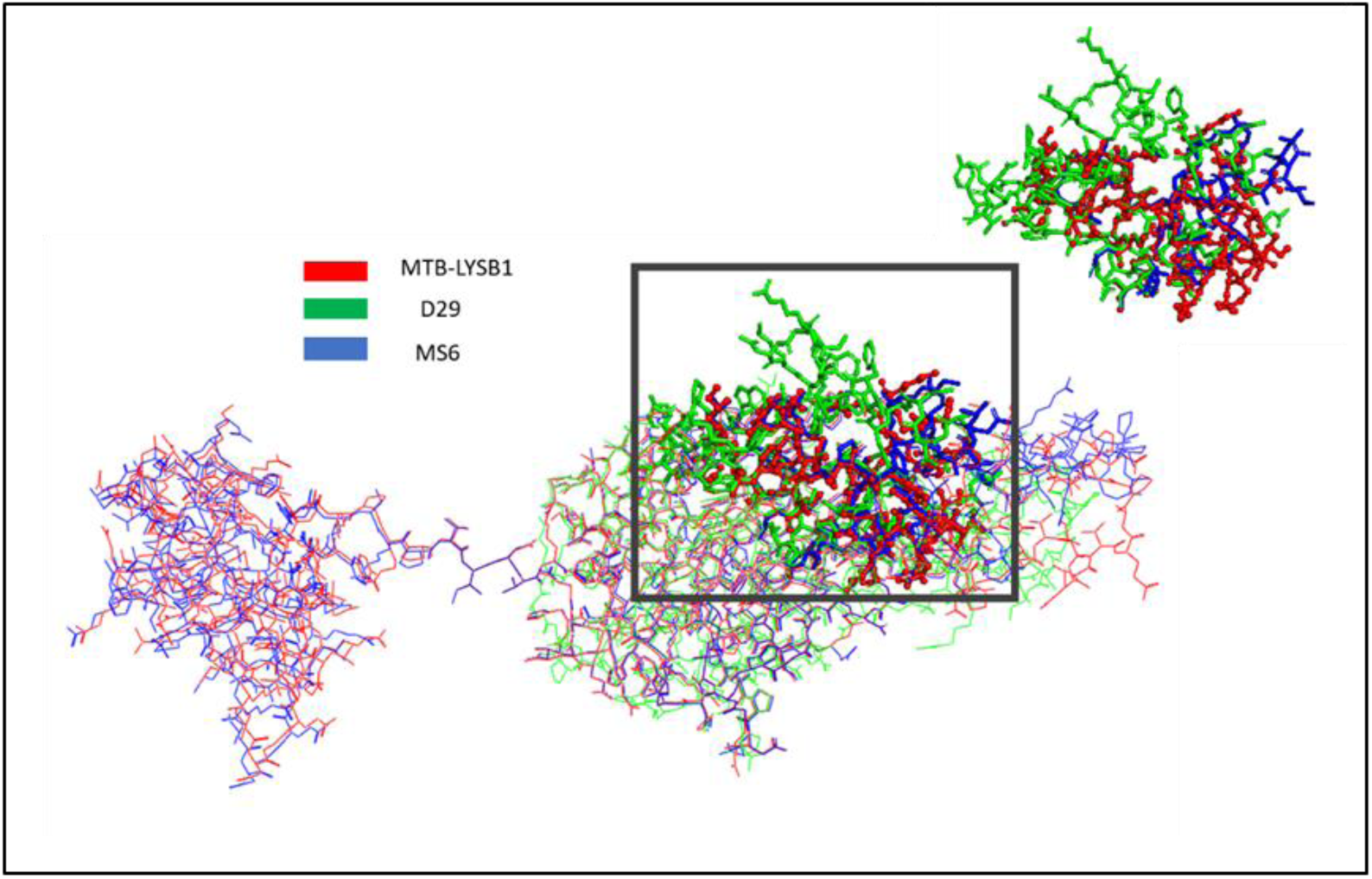
Alignment of binding cavity regions in MTB-LysB1, D29LysB, and Ms6 LysB proteins: Structural superimposition of the three LysB homologues reveals alignment of their predicted binding cavities, identified using CavityPlus. MTB-LysB1 is shown in red, D29 in green, and Ms6 in blue. The inset highlights the cavity-associated region, showing a conserved spatial organisation of the binding pocket across MTB-LysB1 and Ms6, with localised deviations observed in D29.

#### Analysis of Membrane Interaction Motifs

The ester-hydrolysing activity of LysB proteins requires their engagement with the mycobacterial membraneto access the target bonds [20,28]. In D29 LysB, an additional four-helix domain adjacent to its active site has been implicated in lipid substrate binding [20], supporting the notion that surface-exposed hydrophobic features facilitate interaction with membrane lipid components. Furthermore, interfacial enzymes that act on lipid substrates commonly utilise amphipathic helices to adsorb to lipid surfaces, stabilising the enzymes at the membrane and enhancing catalysis [36]. Using HeliQuest, we examined the D29, Ms6, and MTB-LysB1 protein sequences to predict hydrophobic and amphipathic helices, and mapped them onto the modelled structures to assess their membrane association potential (Supplementary Figure S3). While the length and the composition of the predicted helices (Table 2) varied, they were consistently localised near the binding cavity region in all three LysBs, indicating a conserved orientation relative to the membrane, which may play functional roles in membrane targeting, substrate access, and stabilisation of enzyme–substrate interactions during membrane-associated hydrolysis.

**Table 2.** Comparative analysis of hydrophobicity and membrane-interaction features of LysB proteins. Hydrophobicity (H), residue composition, net charge, and hydrophobic face characteristics were analysed for MTB-LysB1, D29, and Ms6 LysBs. The table highlights differences in hydrophobic features such as amino acid composition and hydrophobic faces, which we propose may influence membrane association, substrate interaction, and enzymatic orientation at the lipid interface.

| Features | MTB-LysB1 | D29 LysB | Ms6 LysB |
| --- | --- | --- | --- |
| **Hydrophobicity**<br>**(H)** | 0.663<br>(hydrophobic) | 0.691<br>(hydrophobic) | 0.184<br>(very hydrophilic) |
| **Composition** | 28% polar<br>72% non-polar | 35% polar<br>65% non-polar | 63% polar<br>37% non-polar |
| **Hydrophobic**<br>**moment (μH)** | 0.172<br>(Low amphipathicity) | 0.335<br>(Moderate amphipathicity) | 0.476<br>(Strong amphipathicity) |
| **Net charge (z)** | −1<br>(acidic) | −1<br>(acidic) | +1<br>(basic) |
| **Hydrophobic face** | A-I-F-L-A<br>(Short) | A-L-F-I-G-L-A-I-A-I-I<br>(Long, continuous) | W-A-I-I-A-I<br>(Short) |

### Purification of recombinant MTB-LysB1 enzyme

The MTB-LysB1 gene was cloned into the pET28a expression vector and expressed in *E. coli* BL21 (DE3). The protein obtained from the soluble fractions was purified by affinity chromatography, analysed by SDS-PAGE (Figure 7A) and western blotting, using anti-His antibodies (Figure 7B).

**Figure 7.**
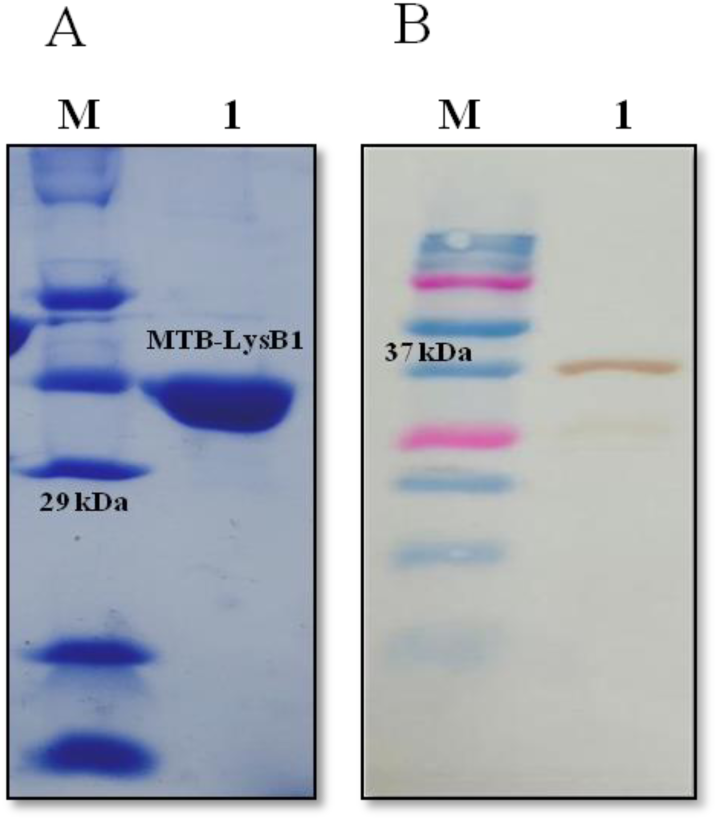
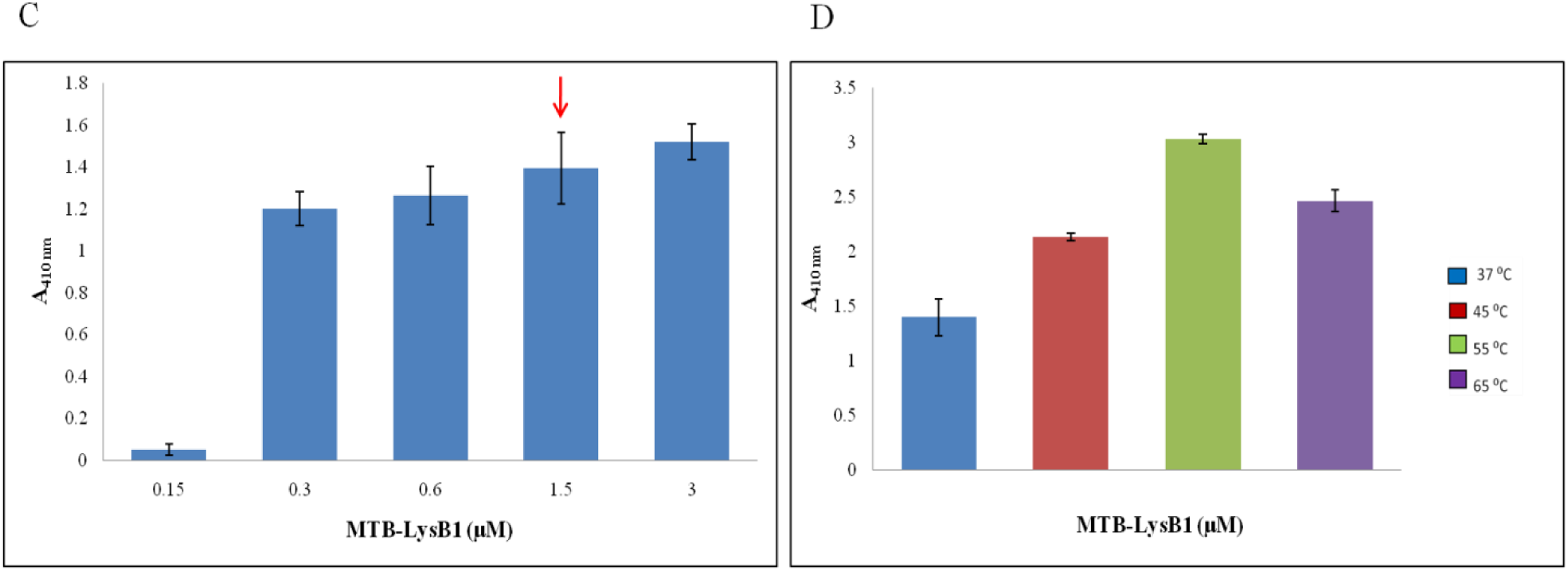
Analysis of purified recombinant MTB-LysB1. **(A)** 12% SDS-PAGE of purified LysB protein stained with Coomassie Brilliant Blue R-250. (**B)** Western blot was developed using 3,3’-diaminobenzidine (DAB) and H_2_O_2_. **Lane M:** Protein molecular weight marker; **Lane 1:** Purified MTB-LysB1 (39 kDa). **(C & D)** pNP-release assay and thermal stability of MTB-LysB1. **(C)** Effect of enzyme concentrations on pNP-release. Reactions were performed at different enzyme concentrations using 10 mM pNPB as the substrate in 25 mM Tris buffer (pH 7.2). The incubation was kept at 37°C for 15 min, and absorbance was measured at 410 nm. The red arrow denotes the enzyme concentration selected for subsequent thermal stability assays. (**D)** Effect of temperature on enzyme activity. Esterase assays were performed at 37°C, 45°C, 55°C and 65°C. All experiments were conducted as two independent sets (n = 4).

### *In vitro* esterase activity of purified MTB-LysB1 enzyme, using pNP-release assay

The esterase activity of purified MTB-LysB1 was evaluated at concentrations ranging from 0.15 to 3 μM using pNPB as the substrate. A sharp increase in activity was observed from 0.15 μM to 0.3 μM, followed by a moderate increase at higher concentrations (Figure 7C). The enzyme’s specific activity was determined to be 5.1 U/mg.

Subsequently, the effect of temperature on enzyme activity was measured to determine its thermal stability, where the activity was observed to increase progressively up to 55°C, with a decline at 65°C, while retaining 83.3% activity relative to 55°C (Figure 7D). Others have also reported a similar thermal stability profile for LysB proteins from D29, Saal, Omega and Obama12 phages [22], where D29 and Obama12 LysBs retained 80% of their activity at 60°C. Thisthermal tolerance mightaccountforconformational stability,maintainingthe enzyme’s catalytic orientation in a complex environment such as the waxy mycolic acid layer ofthemycobacterial cell envelope.

### MTB-LysB1 lytic activity against *M. smegmatis*

The antimycobacterial potential of MTB-LysB1 was first evaluated qualitatively using the plate lysis method and the growth-inhibitory effect against planktonic *M. smegmatis* Mc² 155. The results revealed a clear lysis zone and a decline in bacterial growth, confirming the enzyme’s ability to permeate the mycobacterial cell wall and exert lytic activity (Figure 8A). To visualise the effect on cell morphology, MTB-LysB1-treated *M. smegmatis* cells were analysed by FESEM. Cells appeared clumped and morphologically disrupted compared to the untreated controls, indicating damage to the cell membrane (Figure 8B). We noted a similar effect in our previous study on RitSun LysB [12].

**Figure 8.**
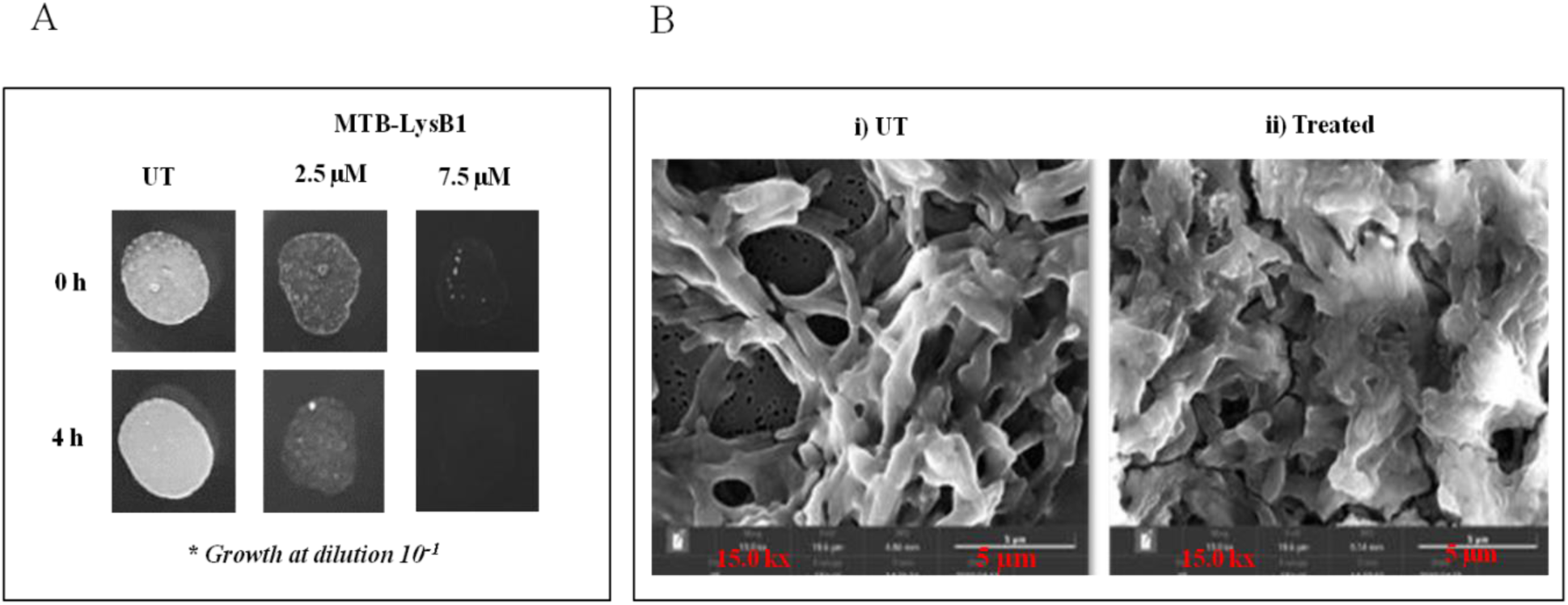
Antibacterial activity and morphological analysis of MTB-LysB1-treated *M. smegmatis* Mc² 155. **(A)** Growth inhibitory effect. Log-phase *M. smegmatis* cells were treated with 2.5 μM & 7.5 μM of MTB-LysB1. Growth inhibition was assessed following a 4 h incubation at 37°C. **(B)** Field Emission Scanning Electron Microscopy (FESEM) analysis. Representative micrographs of *M. smegmatis* Mc² 155 planktonic cells; i) UT (Untreated control), ii) treated with 2.5 μM MTB-LysB1 for 4 h at 37°C. Images were acquired using a TESCAN system at a magnification of 15,000×.

Given the highly hydrophobic nature of the mycobacterial cell envelope and the observed natural permeation of MTB-LysB1, we examined its predicted structure to identify features, such as hydrophobic helices, that might confer cell permeability to our enzyme. We compared it with the D29 LysB structure, which also naturally permeates mycobacterial species (Table 2). Our analysis revealed a high content of hydrophobic residues in both MTB-LysB1 and D29 LysB. This shared hydrophobicity and the predicted hydrophobic helices likely facilitate the ability of these enzymes, both D29 LysB [11] and MTB-LysB1 (this study), to achieve mycobacterial lysis upon exogenous application.

#### Turbidity Reduction Method (TRM) & Log-kill Assay

Further, to quantitatively assess the anti-*M. smegmatis* effect of MTB-LysB1, we performed the Turbidity Reduction Method (TRM) and Log-kill assay. A 67% reduction in OD_600_ relative to the untreated control (Figure 9A) and a notable 3-log reduction in bacterial viability after 24 h of incubation with enzyme (1 μM) (Figure 9B) were observed. At higher concentrations under the tested conditions, a marginal increase ingrowth reduction wasobserved. In the PBS-treated negative control groups, no growth inhibition was observed, confirming that the lytic effect was strictly enzyme-dependent.

**Figure 9.**
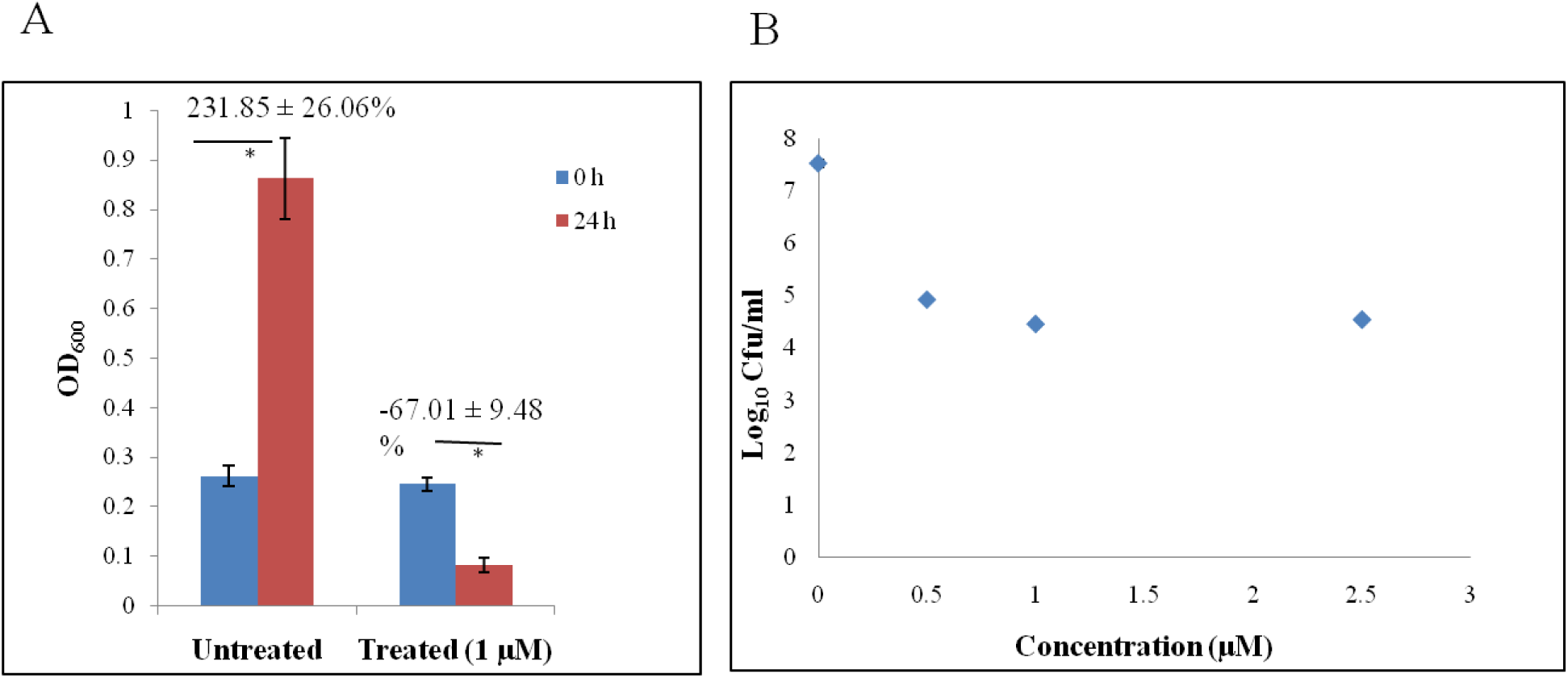
**(A)** Turbidity Reduction Method (TRM). The growth inhibitory effect of MTB-LysB1 treatment on *M. smegmatis* Mc^2^ 155 was assessed by measuring OD_600_ at 0 and 24 h. Growth reduction was calculated relative to the untreated control group, which was normalised to 100% at each time point. All experiments were performed in triplicate (n=3). **(B)** Log Kill Assay. To determine bactericidal activity, *M. smegmatis* cells were treated with varying concentrations of MTB-LysB1 (0.5, 1.0, and 2.0 μM) and incubated at 37°C for 24 h, followed by plating cells on 7H10 agar to enumerate viable colonies. Results are expressed on a logarithmic scale.

### MTB-LysB1 activity against *Mycobacterium tuberculosis*

While assessing MTB-LysB1’s activity against the pathogenic H37Rv strain and a multidrug-resistant *M .tuberculosis* isolate, we observed a zone of clearance in both strains (Supplementary Figure S4), highlighting its host range and clinical relevance.

### Resazurin Microtiter Assay (REMA)

#### M. tuberculosis H37Rv

The antimycobacterial activity of MTB-LysB1 against *M. tuberculosis* H37Rv was evaluated using the REMA assay.We found that the enzyme demonstrated an inhibitory effect (inhibition of resazurin colour conversion from blue to pink) at low micromolar concentrations. Based on visual assessment across independent experiments, the anti-*M. tuberculosis* activities of D29 LysB and MTB-LysB1 were comparable; the approximate MICs of D29 and MTB-LysB were 0.01 µM and 0.01-0.02 µM, respectively. (Figure 10A and 11A). The untreated control wells showed full colour conversion, indicative of active bacterial metabolism.

**Figure 10.**
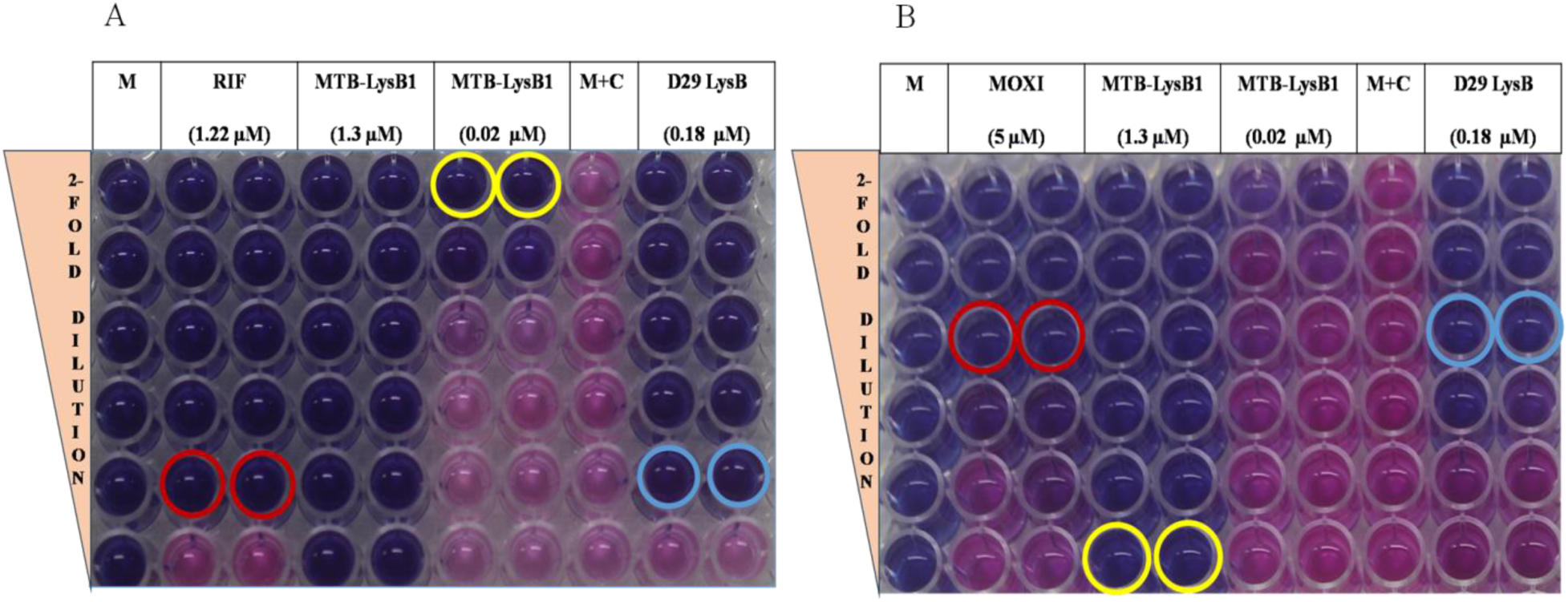
Resazurin Microtiter Assay (REMA). **(A)** Effect of MTB-LysB1 on *M. tuberculosis* H37Rv **B)** Effect of MTB-LysB1 on an MDR isolate. The enzyme demonstrates a concentration- dependent inhibition of metabolic activity against both strains. The retention of the blue resazurin dye indicates growth inhibition, whereas a shift to pink denotes active bacterial growth. **Controls: M** (Media only); **M+C** (Media+Bacterial cells/Growth control). RIF: Rifampicin; MOXI: Moxifloxacin. Circles denote MIC: Yellow (MTB-LysB1), Cyan (D29 LysB), Red [TB Drugs-RIF (A) & MOXI (B)].

**Figure 11.**
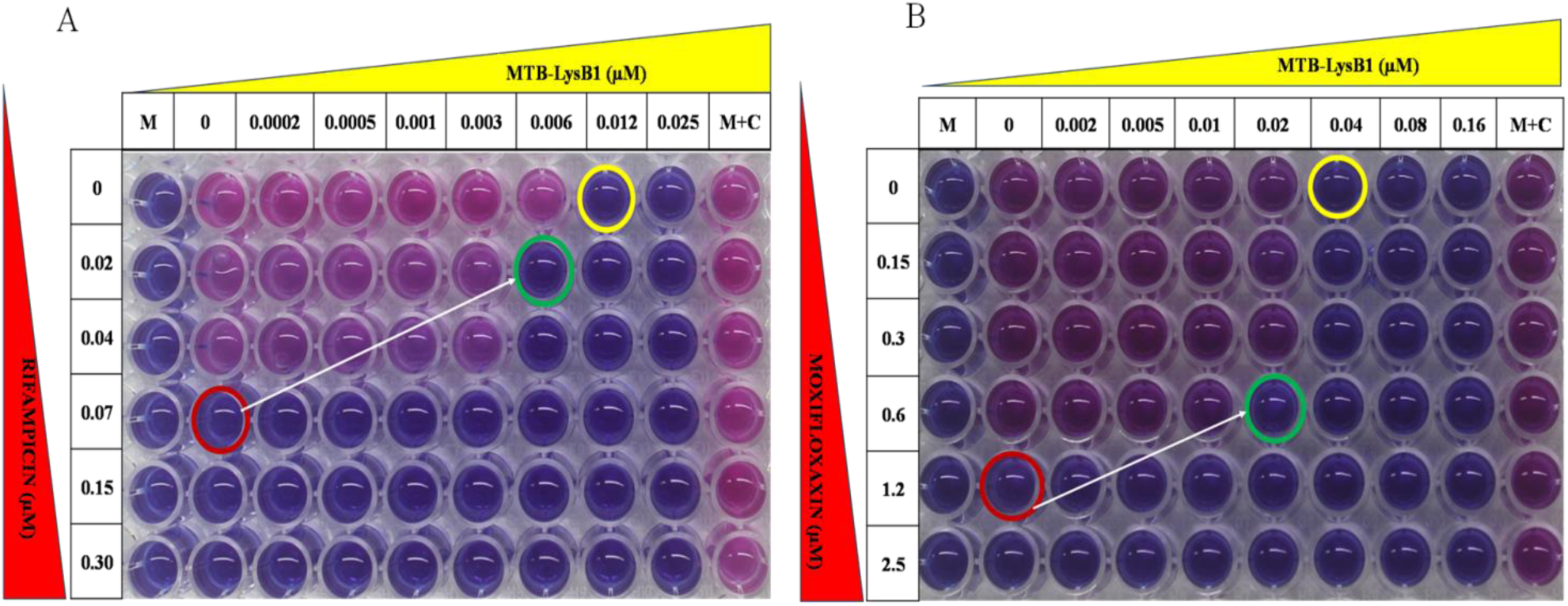
Checkerboard assay to evaluate the combined effects of LysB and TB drugs against *M. tuberculosis* strains. **(A)** Drug-susceptible *M. tuberculosis* H37Rv: MTB-LysB1 and rifampicin were tested using two-fold serial dilutions ranging from 0.025–0.0002 µM and 0.30– 0.02 µM, respectively. **(B)** MDR *M. tuberculosis* isolate: MTB-LysB1 and moxifloxacin were tested using two-fold serial dilutions ranging from 0.16–0.002 µM and 2.5–0.15 µM, respectively. Bacterial viability was determined via REMA. **Controls: M** (Media control); **M+C** (Media + Bacterial cells/Growth control). Circles denote MIC: Yellow (MTB-LysB1), Red [TB Drugs -RIF (A) & MOXI (B)], Green (MTB-LysB1 + TB Drugs)

#### Multi-Drug-Resistant (MDR) M. tuberculosis

On a promising note, the MTB-LysB1 enzyme retained inhibitory activity against a multidrug-resistant (MDR) isolate that was resistant to rifampicin and moxifloxacin (Figure 10B). However, higher enzyme concentrations were required to prevent resazurin reduction compared with the drug-sensitive H37Rv strain; complete inhibition of colour change was obtained at an approximate MIC of 0.04 µM. This shift in susceptibility highlights strain-dependent differences, resulting in a higher threshold for inhibition compared to the standard laboratory strain, while indicating that conventional drug resistance mechanisms do not compromise MTB-LysB1 activity.

### Checkerboard Assay

Next, we performed the checkerboard assay to study the effect of MTB-LysB1 in combination with the TB drug rifampicin fordrug-susceptible *M. tuberculosis* H37Rv and with moxifloxacin for the MDR isolate. We found that the MIC of rifampicin decreased from 0.07 to 0.02 µM when combined with MTB-LysB1, and that of MTB-LysB1 decreased from 0.012 µM to 0.006 µM (Figure 11A). For the MDR isolate, the MICs of moxifloxacin and MTB-LysB1 were lowered from 1.2 to 0.6 µM and from 0.04to 0.02 µM, respectively (Figure 11B). Forboth*M. tuberculosis* H37Rv (rifampicin & MTB-LysB1) and the MDR isolate (moxifloxacin & MTB-LysB1), an additive effect was observed (Table 3).

**Table 3.** Checkerboard assay of MTB-LysB1 in combination with rifampicin against drug-susceptible *M. tuberculosis* H37Rv strain and with moxifloxacin against MDR *M. tuberculosis* isolate, respectively. FICI (Fractional Inhibitory Concentration Index)*.

| *M. tuberculosis* Strain | MTB-LysB1 MIC ($\mu\text{M}$) | DrugMIC ($\mu\text{M}$) | MTB-LysB1 MIC in combination with drug($\mu\text{M}$) | Drug MIC in combination with MTB-LysB1($\mu\text{M}$) | FICI | Interpretation |
| --- | --- | --- | --- | --- | --- | --- |
| H37Rv | 0.012 | RIF: 0.07 | 0.006 | RIF: 0.02 | 0.79 | Additive |
| MDR | 0.04 | MOXI: 1.2 | 0.02 | MOXI:0.6 | 1 | Additive |

\*Additive: $0.5 < \text{FICI} \leq 1.0$ ; Synergy: $\text{FICI} \leq 0.5$ , Indifferent: $1.0 < \text{FICI} \leq 4.0$ , and Antagonism: $\text{FICI} > 4.0$

## Conclusion

The emergence of multidrug-resistant tuberculosis (MDR-TB) continues to challenge current treatment strategies and emphasises an urgent need for therapeutics with novel mechanisms of action. Bacteriophages and the lytic enzymes they encode are gaining significant attention as biotherapeutics for drug-resistant infections [11,37,38]. Bacteriophage-derived endolysins, characterised by rapid bactericidal action and a remarkably low propensity to develop resistance [1,2,8], offer a transformative approach to treating drug-resistant pathogens.In this study, we identified and comprehensively characterised MTB-LysB1, a novel mycobacteriophage-derived LysB, and demonstrated its potential as an adjunctive anti-tubercular therapeutic.

Through integrated structural analyses, we note that despite sequence divergence from previously characterised LysB homologues, MTB-LysB1 carries the conserved α/β-hydrolase fold and catalytic architecture characteristic of the LysB family. Molecular dynamics simulations of MTB-LysB1 confirmed the stability of the predicted structure, and comparative analysesidentified conserved hydrophobic motifs that provide a plausible structural basis for membrane interaction, contributing to its natural cell-permeabilityand enzyme activity.

Biochemical characterisation confirmed that recombinant MTB-LysB1 is an active and thermostable esterase. The enzyme exhibited potent lytic activity against *M. smegmatis*, producing extensive cell-envelope disruption and a 3-log reduction in bacterial viability. More importantly, MTB-LysB1 demonstrated potent anti-tubercular activity against both drug-susceptible *M. tuberculosis* H37Rv and a multidrug-resistant clinical isolate, with minimum inhibitory concentrations in the low micromolar range. In combination with rifampicin or moxifloxacin, MTB-LysB1 produced additive effects, reducing the effective concentrations of both the enzyme and the companion antibiotics, highlighting its potential to improve treatment efficacy while reducing antibiotic dose.

Collectively, this study identifies MTB-LysB1 as a structurally stable and biologically active anti-tubercular candidate that warrants further preclinical evaluation as an adjunct to existing tuberculosis therapies. Beyond demonstrating its anti-tubercular efficacy, our work provides structural insights into membrane interaction and catalytic function that may guide the rational engineering of next-generation LysB enzymes. With improved understanding of biopharmaceutical production and targeted delivery formulations, phage-derived endolysins such as MTB-LysB1 represent promising adjuncts to conventional antitubercular therapy, especially for treating MDR infections and may also offer broader applications against non-tuberculous mycobacterial infections.

## Supplementary Information

**Figure S1:**
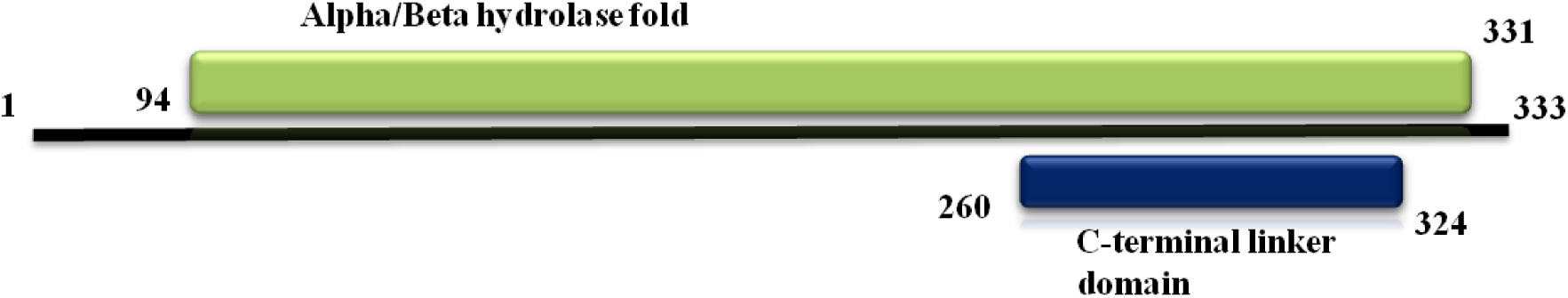
Domain organisation in MTB-LysB1 (333 aa). The InterProScan analysis predicted two domains: an alpha/beta hydrolase fold (94-331 aa) and a C-terminal linker domain (260-324 aa) embedded in the alpha/beta hydrolase fold. The N-terminal region: 1-93 aa.

**Figure S2.**
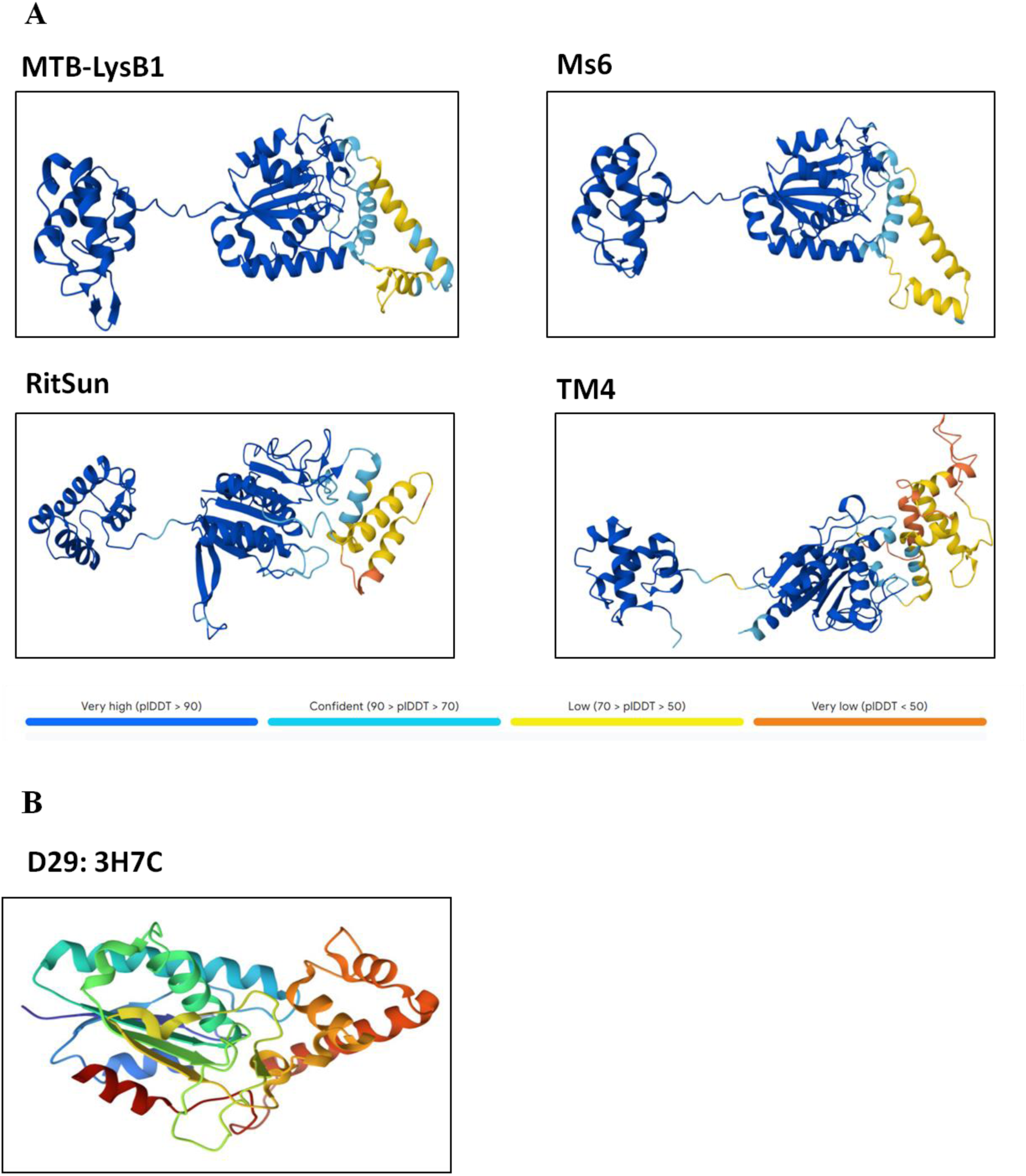
**A)** AlphaFold2 Modelled structures of LysBs represented in cartoon format and coloured according to confidence score. The proteins exhibit a conserved central catalytic domain, predominantly composed of α-helices and β-sheets, connected to variable terminal domains via flexible linker regions. Structural differences are mainly observed in flexible loop and terminal regions. **B)** PDB structure of D29 LysB. PDB ID: 3HC7.

**Figure S3:**
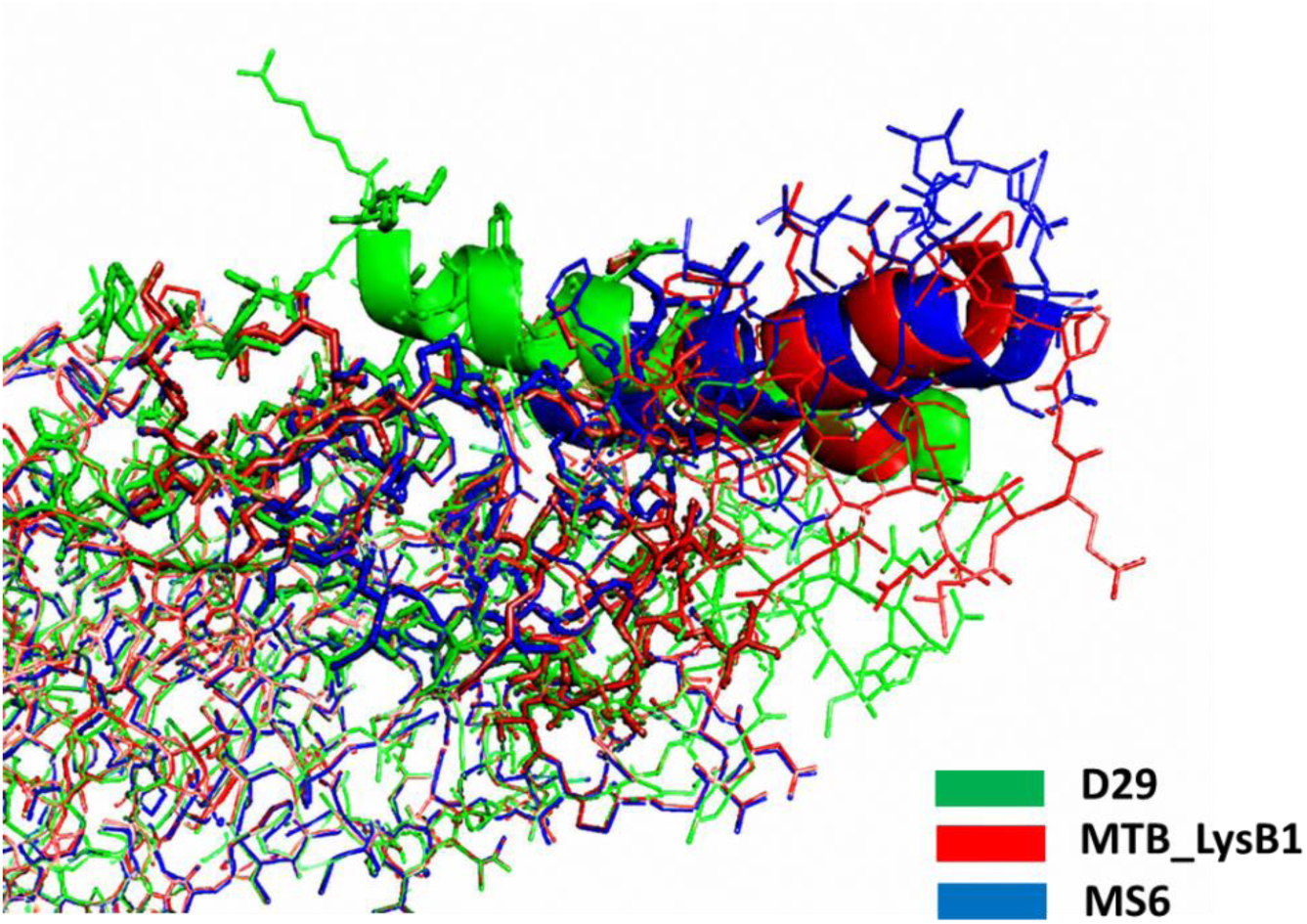
Hydrophobic helix comparison among LysB homologues: Superimposition of the predicted hydrophobic helical regions of MTB-LysB1, D29 LysB, and Ms6 LysB highlights notable differences in physicochemical properties. MTB-LysB1 and D29 LysB exhibit predominantly hydrophobic helices, suggesting stronger membrane-embedding potential, whereas Ms6 LysB displays comparatively hydrophilic helical regions, indicating possible adaptation toward more aqueous environments.

**Figure S4.**
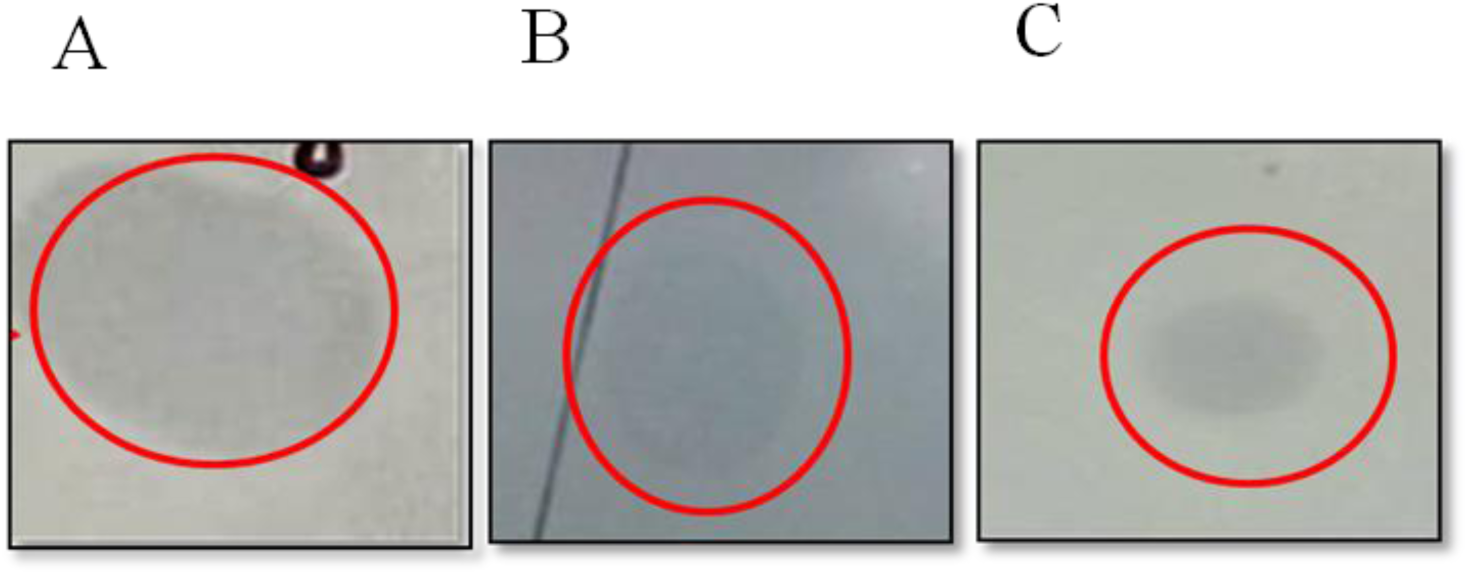
Plate lysis method to assess MTB-LysB1 lytic activity against **(A)** *M. smegmatis,* **(B)** *M. tuberculosis* H37Rv, and **(C)** *M. tuberculosis* MDR strain.

## Acknowledgements

We thank Anusandhan National Research Foundation (ANRF), previously Science and Engineering Research Board (SERB), GoI, New Delhi, India, for supporting the project (EMR/2017/004051). **RA** received Senior Research Fellowship (SRF) from the University Grants Commission (UGC), New Delhi, India. **EK** received TCOF fellowship from the Council of Scientific and Industrial Research (CSIR). **JR** received Senior Research Fellowship (SRF) from the Indian Council of Medical Research (ICMR), New Delhi, India. We thank Acharya Narendra Dev College (ANDC), University of Delhi, New Delhi, India, and the Experimental Animal Facility at NJIL&OMD, Agra,India, for providing the infrastructural support. We used Google Gemini in the manuscript preparation.

## Credit authorship contribution statement

**RA:** Writing–original draft & editing, Methodology, Experiments, Data curation, Validation. **EK:** Writing–review & editing, Methodology, Experiments. **JR:** Writing–original draft & editing, Methodology, Data curation, Visualisation. **AKS:** Writing–original draft, Methodology, Experiments. **UB:** Conceptualisation, Funding acquisition, Writing–review & editing, Validation, Supervision, Project administration.All the authors have read and approved the final manuscript.

